# Production of Mixed Fruits (Watermelon, Banana, and Pineapple) Wine Using *Saccharomyces cerevisiae* Isolated from Palm Wine

**DOI:** 10.64898/2026.02.28.708690

**Authors:** Uwamere Edeghor, Jessica Egbelo, Jonah Nwokpuru, Chioma Achokwu, Victory Igwe

## Abstract

Postharvest losses and rapid nutrient degradation due to fruit spoilage necessitate alternative preservation methods. Wine production presents a viable approach to minimizing fruit waste while retaining essential nutrients. In this study, mixed fruit wines (watermelon, banana, and pineapple) were produced using *Saccharomyces cerevisiae* isolated from palm wine as a starter culture. After secondary fermentation, the wines maintained an acidic pH range (2.29±0.1 to 3.25±0.2), a stable fermentation temperature (26.50±1.1°C to 27.00±1.1°C), specific gravity values of 1.021±0.02 kg/L and 1.027±0.03 kg/L, and total acidity levels of 1.57±0.2% and 2.11±0.1% for Wines A and B, respectively. The final alcohol content was 8.40±2.9% in Wine A and 9.84±3.6% in Wine B. Proximate analysis demonstrated the retention of key nutrients post-clarification and maturation, and sensory evaluation indicated a higher consumer preference for Wine B (*P*>0.05). These findings highlight the potential of indigenous *S. cerevisiae* strains from palm win for efficient wine fermentation and support the utilization of mixed fruits as a sustainable raw material for value-added wine production. This approach not only mitigates fruit wastage but also provides an economic avenue for enhancing fruit utilization.

## 1.0 INTRODUCTION

### DEFINE THE PROBLEM

Most tropical fruits are sensitive to low-temperature storage with an optimum storage temperature of 13 ± 1ºC. Physiological disorders resulting from metabolic responses of the fruit to the storage environment decrease their marketable values (Wongs-Aree and Noochinda, 2014). A lot of fruits are grown in the three zones of the tropics. However, only few are exploited and utilized in local markets and fewer are exported. The majority of the export market is made up of banana (*Musa sapientum*), mango (*Mangifera indica*), pineapple (*Ananas comosus*), citrus, papaya (*Carica papaya*), avocado (*Persea americana*), watermelon (*Citrullus lanatus*) although some may categorize it as a vegetable depending on how it is being consumed. Tropical fruits, in most cases, are sold and eaten fresh (except breadfruit, nutmeg and sibu olive), while off-grade fruit is processed. Very often tropical fruits are turned into juices, chips, purees, jellies, pickles, chutneys, flour, wines and enzymes (end result of fermented fruits which is believed to be good for the gut) (Ding, 2017). Tropical countries like Nigeria cultivate different fruits; some are termed rare, exotic or tropical fruits. Due to their high sugar content and nutritional value, they are good for wine production. Some other parts of the fruits are also used especially in Nigeria, such as the seed, fruit skin, leaves and even the tree bark, which serve for medicinal purposes, source of raw materials and food to both man and animal.

### LITERATURE REVIEW

Fruits make up a large portion of our diets. The botanical definition of fruit is a seed-bearing part of a flowering plant or tree that can be eaten as foods, such as avocados, cucumber, squash and yes, even tomatoes are all fruits. From a culinary viewpoint, a fruit is usually thought of as any sweet tasting point product with seeds (Bialowas and Ricketts, 2021). Tropical countries like Nigeria cultivate different fruits; some are termed rare, exotic or tropical fruits. Due to their high sugar content and nutritional value they are good for wine production. Some other parts of the fruits are also used especially in Nigeria, such as the seed, fruit skin, leaves and even the tree bark, which serve for medicinal purposes, source of raw materials and food to both man and animal. Very often tropical fruits are turned into juices, chips, purees, jellies, pickles, chutneys, flour, wines and enzymes (end result of fermented fruits which is believed to be good for the gut) (Ding, 2017). Wine can be made from virtually many plant matters that can be fermented (Harding, 2005). Wine making involves the use of yeast to ferment the must of a chosen fruit or fruits for a number of days, depending on the objective of the winemaker. The yeast which is the main organism responsible for alcoholic fermentation usually belongs to the genus *Saccharomyces* (Okeke *et al*., 2015). Yeasts from other sources such as palm wine have also been used in the production of fruit wine (Ayogu, 1999).

Watermelon (*Citrullus lanatus*) is an exotic quintessential fruit that contains nutrients and phytochemicals reported to be beneficial to human health (Choudhary *et al*., 2015; Ijah *et al*., 2015). It is a good source of vitamins B, C, and E as well as minerals such as phosphorus, magnesium, calcium, and iron (Romdhane *et al*., 2017). Epidemiological studies have demonstrated that it possesses antioxidants with anti-inflammatory, antihypertensive properties as well as a protective effect against carbon tetrachloride-induced toxicity (Choudhary *et al*., 2015; Ijah *et al*., 2015). Consumption of raw watermelon fruit on hot summer days is a common practice which has been observed across the world; however, to increase utilization and availability throughout the year, watermelon is processed into variety of commercial products (Said, 2014). It has been used in the production of a variety of products like juice, smoothies, jams, sweets, and sauces (Jumde *et al*., 2015). Watermelon fruits yield 55.3% juice, 31.5% rind, and 10.4% pomace. The sweetness of watermelon is mainly due to the combination of sucrose, glucose and fructose. Sucrose and glucose account for 20-40% and fructose for 30-50% of total sugars in a ripe watermelon (Bianchi *et al*., 2018).

Banana (*Musa sapientum*) is an important staple starchy food in Nigeria. Ripe bananas are consumed raw as a desert fruit. Banana serves as good nutritional sources of carbohydrates, minerals such as potassium and vitamins such as B1, B2, B3, B12, C and E. Following the high nutritional content of banana, it is consumed in large quantity in a variety of ways in Africa. The banana fruit can be eaten raw or cooked, processed into flour or fermented for the production of beverages such as banana juice, beer, vinegar and wine (Pillay *et al*., 2004; Pillay and Tripathi, 2007). However, banana has a short shelf-life under the prevailing temperature and humidity condition in tropical countries, including Nigeria. This results to wastage of the fruits as a result of poor handling and inadequate storage facilities (Akubor *et al*., 2003; Wall, 2006). Moreover, fermenting banana juice into wine is considered to be an attractive means of utilizing surplus banana, since the consumption of banana wine provides a rich source of vitamins and ensures harnessing of the fruits into a useful by-product (Obaedo and Ikenebomeh, 2009).

Pineapple (*Ananas comosus*) is a fruit which has a dense texture is rich in vitamins, enzymes, and anti-oxidants (Hossain *et al*., 2015). It is a perennial monocotyledonous herb, with a short basal stem, producing adventitious roots below and a crown of spirally arranged leaves. It produces a single syncarpous fruit on a terminal inflorescence. The fruit is juicy with excellent flavor and taste. Pineapple is now considered to be the third most important fruit crop in world production after banana and citrus (Malezieux *et al*., 2003). Pineapple contains a proteolytic enzyme, bromelain which helps in the digestion process by breaking down proteins. Bromelain has anti-inflammatory, anti-clotting and anti-cancer properties. Furthermore, it can also interact with other medications (Amini *et al*., 2014).

Wine is any alcoholic beverage produced from juices of variety of fruits by fermentative action of microorganisms either spontaneously or seeding with a particular strain mainly of yeast species to adopt a particular quality of wine (Okeke *et al*., 2015). Wine is a popular alcoholic drink and the process of wine preparation is known as vinification, the branch of science that deals with the study of wine is known as enology (Gaurab, 2020) and it plays an important role in meals but also in the protective effects against cardiovascular disease, it is a complex mixture of different compounds at several concentrations (Ferrer-Gallego, 2011).

Wine has been made from the fermentation of grapes for nearly 7000 years now (Adams *et al*., 2004). The process of fermenting is basically feeding sugars and nutrients in solution to yeast, which return the favor by producing carbon dioxide gas and alcohol. This process goes on until either all the sugar is gone or the yeast can no longer tolerate the alcoholic percentage of the beverage. Yeast has the capability of converting grapes into an alcoholic compound and removing the sugar content in it for the production of different types of wines (Okeke *et al*., 2015). The yeast to carry out the fermentation process is isolated from the Nigerian alcoholic beverage known as palm wine.

Palm wine is the fermented sap of the tropical plant of the *palmae* family. It is produced and consumed in very large quantities in the Southeastern Nigeria. It contains nutritionally important components including amino acids, proteins, vitamins and sugar (Okafor, 2007). Palm wine is presented in a variety of flavors, ranging from sweet (unfermented) to sour (fermented) and vinegary. It is collected by tapping the top of the trunk; it is a cloudy, whitish beverage with a sweet alcoholic taste and a very short shelf life of only 1 day. It is produced by a succession of microorganisms, Gram-negative bacteria, lactic acid bacteria and yeasts as well as acetic acid bacteria. Yeasts isolated from palm wine have been identified as coming from various genera such as *Saccharomyces, Pichia, Schizosaccharomyces, Kloekera, Endomycopsis, Saccharomzeoides* and *Candida* which find their way into the wine from a variety of sources including air, tapping utensils, previous brew and the trees. Hence, palm wine serves as a source of single cell protein and vitamins (Fleet 2003; Ezereonye 2004; Okafor 2007; Duarte *et al*. 2010; Adedayo *et al*., 2011).

Winemakers consider several factors during the production of wine of which sugar content of the juice and yeast strain deployed during the fermentation process are paramount (Okemini and Dilim, 2017). Generally, consumer’s preference for a particular brand and/or type of wine is influenced by the aroma, color, taste, quality, guaranteed origin, ecological production and other perceived sensory properties of the product (Saranraj *et al*., 2017).

### NOVELTY OF THE RESEARCH

The objective of this study is to produce quality wine from mixed fruits: watermelon, banana and pineapple using the isolated yeast strain, *Saccharomyces cerevisiae* from palm wine.

## 2.0 Methodology

### 2.1 Sample collection

Ripe watermelon (*Citrullus lanatus*), banana (*Musa sapientum*) and pineapple (*Ananas comosus*) were purchased from the local market (Ika Ika Oqua (Marian) Market) in Calabar Municipality, Cross River State, Nigeria. Fresh palm wine was purchased from a palm wine tapper at Atimbo, Calabar Municipality, Cross River State. The fruits and palm wine were transported to the laboratory in clean disposable bags and in an ice box respectively for analysis.

### 2.2 Isolation of *Saccharomyces cerevisiae* from palm wine

Yeasts were isolated using the method described by Abosede *et al*. (2013). One milliliter of freshly tapped palm sap was diluted serially in ten folds using sterile distilled water as diluents. Dilutions of 10^-4^, 10^-5^ and 10^-6^ were cultured in duplicates in Malt Extract Agar (MEA) using pour plate technique. The plates were incubated at 30°C for 48hrs. The yeast cultures were transferred to a modified Malt Extract Agar (MEA) containing 2% glucose and then incubated for 24hrs. Seven out of the ten isolates were identified as *Saccharomyces cerevisiae* based on their cultural characteristics, microscopy and their pattern of fermentation and assimilation of glucose, maltose, sucrose, lactose, saccharose and fructose as described by Amoa-Awua *et al*. (2006). The identified organism Colonies were counted after which the different yeast isolates were purified. Ten isolates were obtained and sub-cultured on fresh medium to obtain pure cultures. was maintained on MEA slant.

### 2.3 Preparation of must for mixed fruit fermentation

The must was prepared for two mixed fruit fermentations: watermelon and pineapple (wine A) & watermelon and banana (wine B). The fruits were thoroughly washed with distilled water and then peeled. Exactly 450g each of the fruit samples, watermelon, banana and pineapple were weighed for two-mixed fruit fermentation. The fruits were cut into bits using a clean knife before transferring them as measured into a laboratory blender (Model: BLG-605SS) containing 100ml of sterile water for crushing. The crushed sample was filtered into a clean, new transparent bucket to obtain the must. Exactly 118g of sucrose and 50g of table sugar were added to the must for fortification and stirred vigorously. This was followed by the addition of 0.75g of sodium metabisulfate (Na_2_S_2_O_5_) which serves as a sterilizer and prevents fermentation before the addition of the yeast starter.

### 2.4 Inoculum development

Inoculums’ development was done to obtain large quantity of yeast cells to build up in order to carry out the fermentation. Ten milliliters of sterile water was poured into a test tube, caped and autoclaved at 121°C for 15 minutes. It was allowed to cool and 2 loopfuls of the identified *Saccharomyces cerevisiae* was diluted in 10ml of sterile water, mixed properly and allowed to stand for 1hr. Two milliliters of the prepared inoculums which is a representation of 4.7 x 10^6^ yeast cells, was taken and used to induce fermentation.

### 2.5 Must fermentation

The must was then transferred into a sterile 1000ml conical flask and this was followed by the addition of 2.5g ammonium sulfate, 5g of magnesium sulfate and 5g of potassium dihydrogen for yeast supplementation. The must was inoculated with yeast obtained by inoculum development and was mixed properly. The must was dispensed into five 100ml conical flasks covered with an airlock and allowed to ferment for a period of 14 days. During this period, primary fermentation was monitored for 4 days at 12h interval by carrying out microbiological and physicochemical (pH, temperature, reducing sugar, specific gravity, alcohol content and total acidity) analysis. After 14 days, secondary fermentation was achieved by filtering using a sterilized muslin cloth and siphon tubes. The residue was carefully removed and decanted into another flask, thereafter 9g of granulated sugar was added. Exactly 0.5g of gelatin was dissolved in 50mls of boiling water and stirred properly to a gel form and allowed to stand for 24h. Then 0.75ml of the gel-like gelatin was transferred into each wine followed by stirring to dissolve properly. A small quantity of the mixture was collected and placed in a clean bottle containing the wine which was covered tightly and was used to monitor the process of clarification. This was done for a period of one week. Filtration was done after the wines had completed clarification using a clean cloth and siphon tubes sterilized by using 70% alcohol. The wine residues were removed and the filtrates were allowed to mature and sensory evaluation was done.

### 2.6 Microbiological analysis

Microbial analysis of the wine was carried out using Malt extract agar (MEA) and prepared according to the manufacturer’s instruction. During primary fermentation at 12hrs interval, 1ml of the must was sampled and serially diluted, 0.1ml of the dilute was cultured on Malt extract agar treated with chloramphenicol using spread plate method, incubated for 24hrs and observed for growth. The colonies in the culture plates were enumerated and the microbial population was calculated, the results were expressed in CFU/ml.

### 2.7 Physiochemical analysis

Sampling was carried out during primary fermentation and after secondary fermentation for the determination of pH using a pH meter (Model: PH-9808I), temperature using an analytical thermometer, reducing sugar using the method described by Miller (1971). The determination of specific gravity was done by using a density bottle while total acidity was determined by the methods described by Amerine and Ough (1980). Alcohol content was determined by the difference in specific gravity.

### 2.8 Proximate analysis

Proximate composition covered the determination of sample quantitatively in terms of protein, fat, fiber, ash, moisture content/dry matter, carbohydrate content and energy using the method described by FAO (1988), John Kjeldhal (1883) which was modified by Ibitoye (2008) and Chapman and Pratt (1961).

### 2.9 Organoleptic evaluation

This is the use of human senses for the purpose of evaluating consumer products. It requires panels of human assessors, on whom the responses made by them on the products are recorded. The wines produced were compared for color, flavor, taste, clarity, and overall acceptability by a panel of ten judges on a five-point hedonic scale where five denotes excellent and one very poor.

### 2.10 Statistical analysis

The completely randomized analysis of variance (ANOVA) was used as described by Winner (2004) to analyze the data obtained. Descriptive analysis, mean separation and comparison were done using Microsoft Excel 2010. Significant difference was accepted at P<0.05 and results were expressed as mean ± standard deviation from the mean.

## 3.0 Results

The pH was observed throughout the period of primary fermentation to range from 3.69 to 3.32 in watermelon and pineapple wine (Wine A) and 3.84 to 3.34 in watermelon and banana wine (Wine B) from 0 to 96hrs as shown in Figure 1. Temperature of the wines during fermentation ranged from 30°C to 28°C in Wine A and 29°C to 27.5°C in Wine B from 0 to 96hrs as shown in Figure 2. In the case of reducing sugars, the values were observed to range from 0.0440mg/1000ml to 0.0230mg/1000ml in Wine A and 0.0449mg/1000ml to 0.0340mg/1000ml in Wine B from 0 to 96hrs as depicted in Figure 3. Specific gravity of the wines as displayed in Figure 4, ranged from 1.085kg/l to 1.023kg/l in Wine A and 1.102kg/l to 1.028kg/l in Wine B. In Figure 5, the alcohol content of the wines ranged from 0 to 8.14% in Wine A and 0 to 9.71% in Wine B.

**Fig. i:**
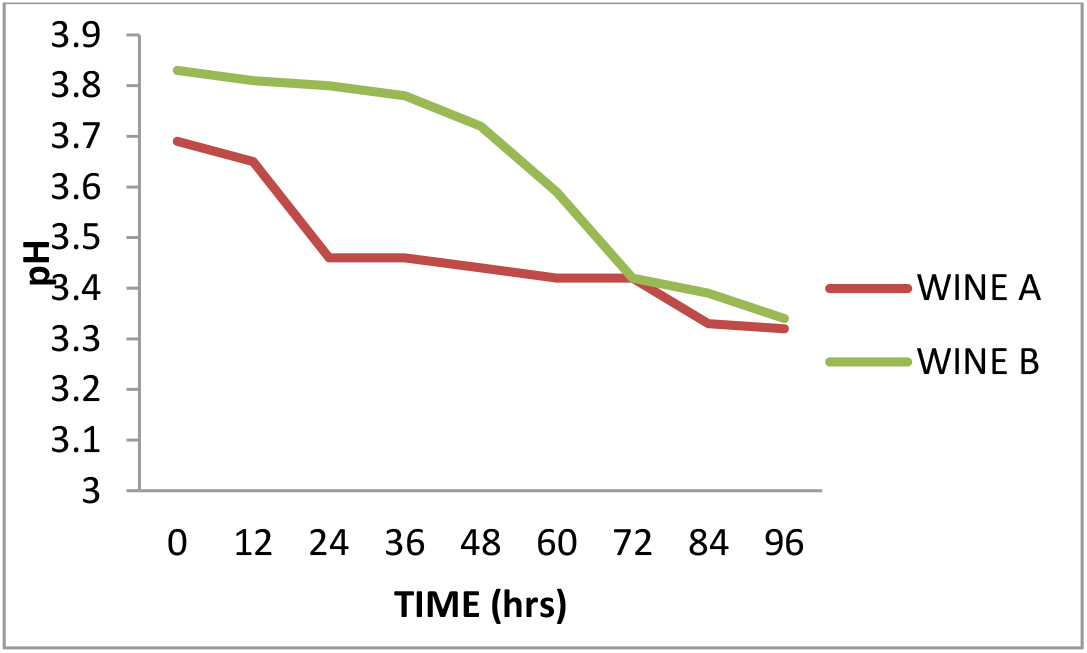
Variation in the pH of the wines during primary fermentation

**Fig. ii:**
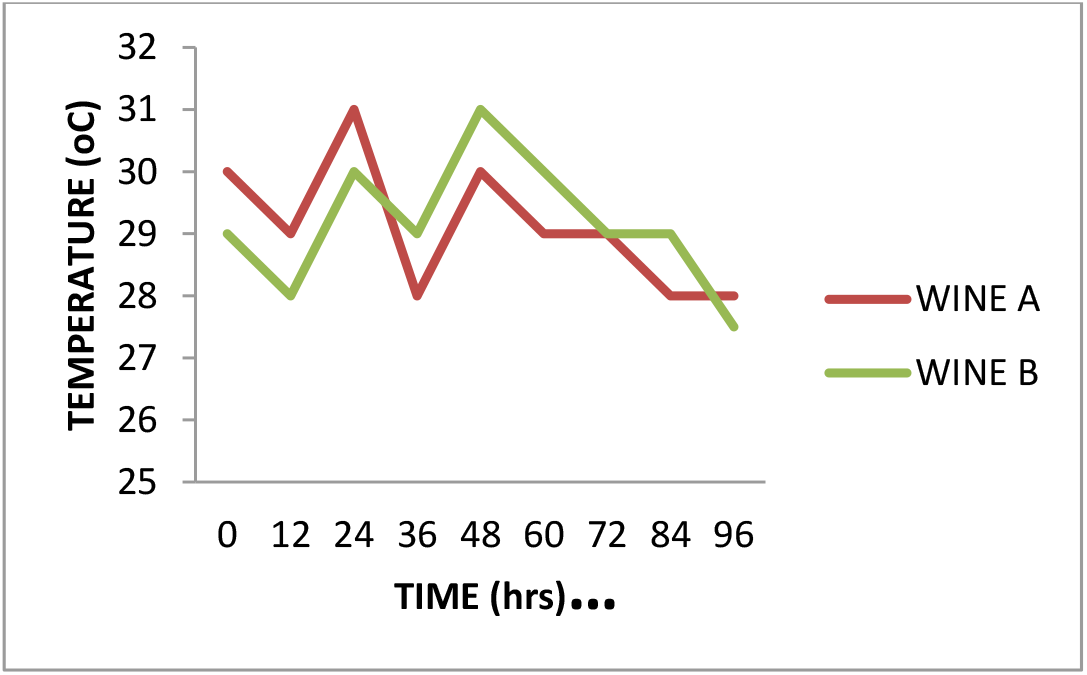
Variation in the temperature of the wines during primary fermentation

**Fig. iii:**
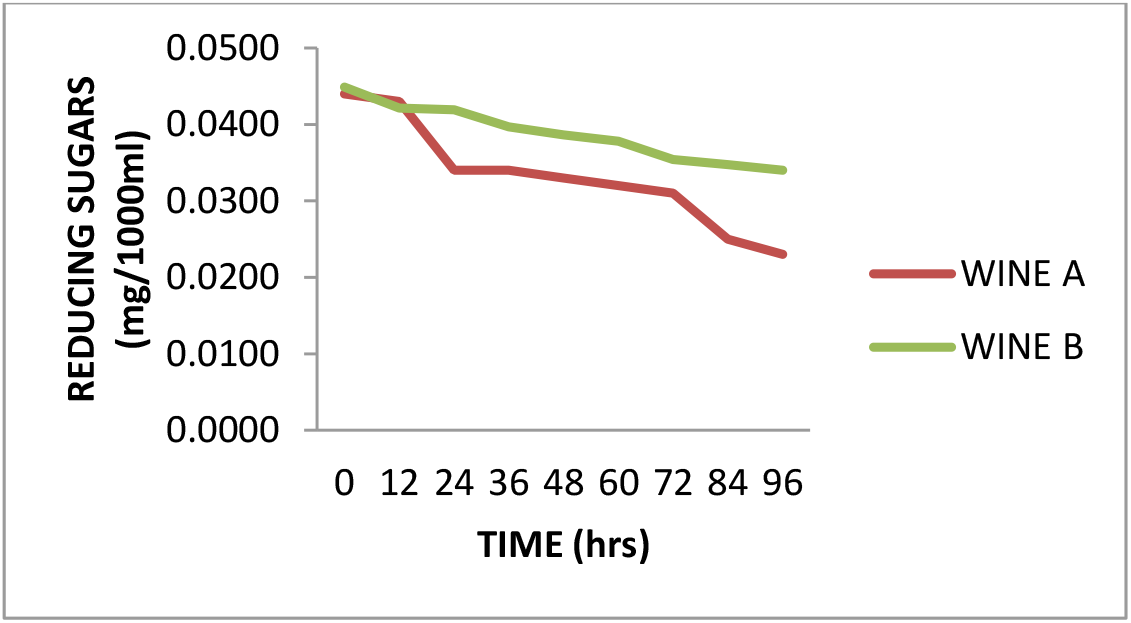
Variation in the reducing sugars of the wines during primary fermentation

**Fig. iv:**
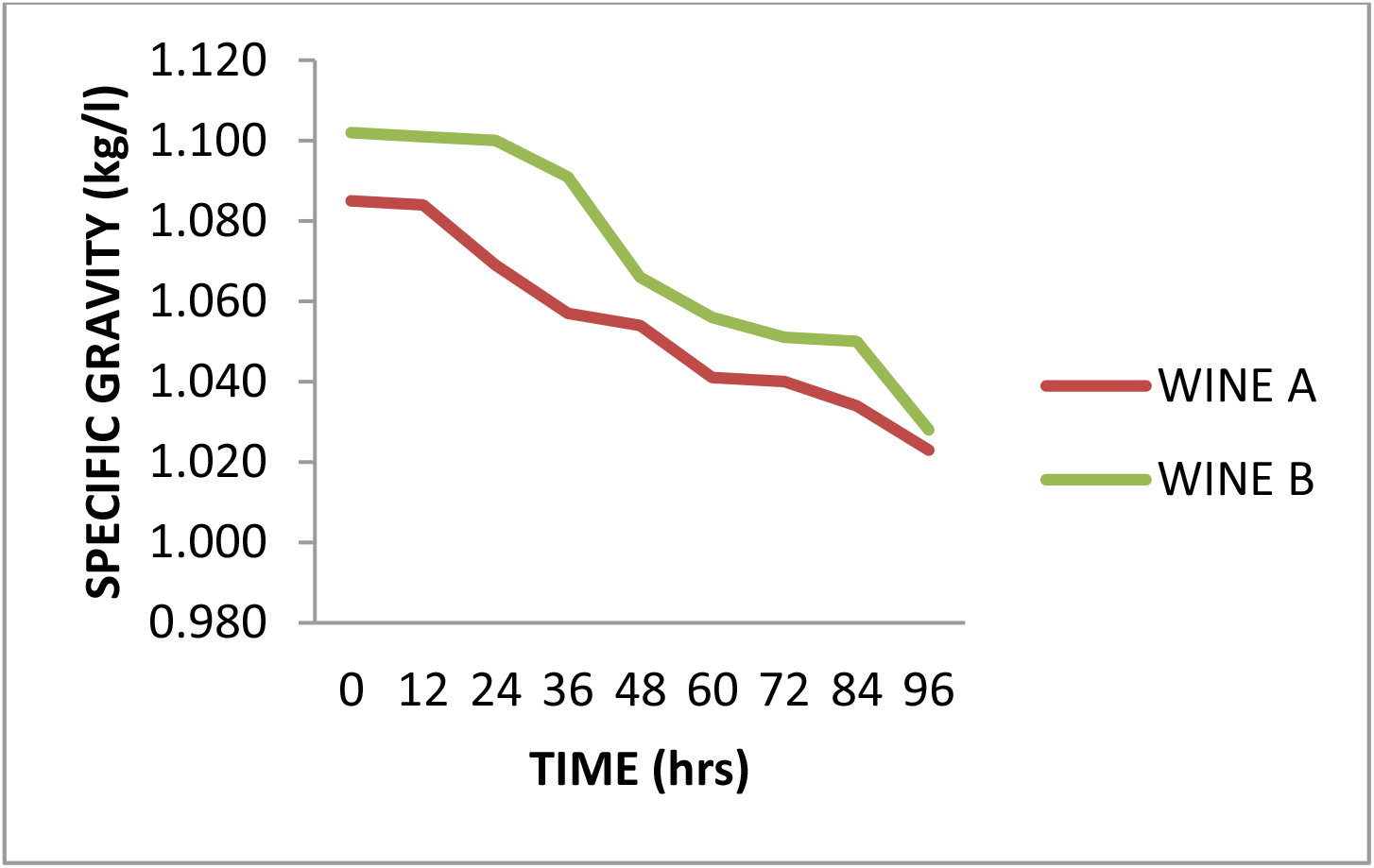
Variation in the specific gravity of the wines during primary fermentation

**Fig. v:**
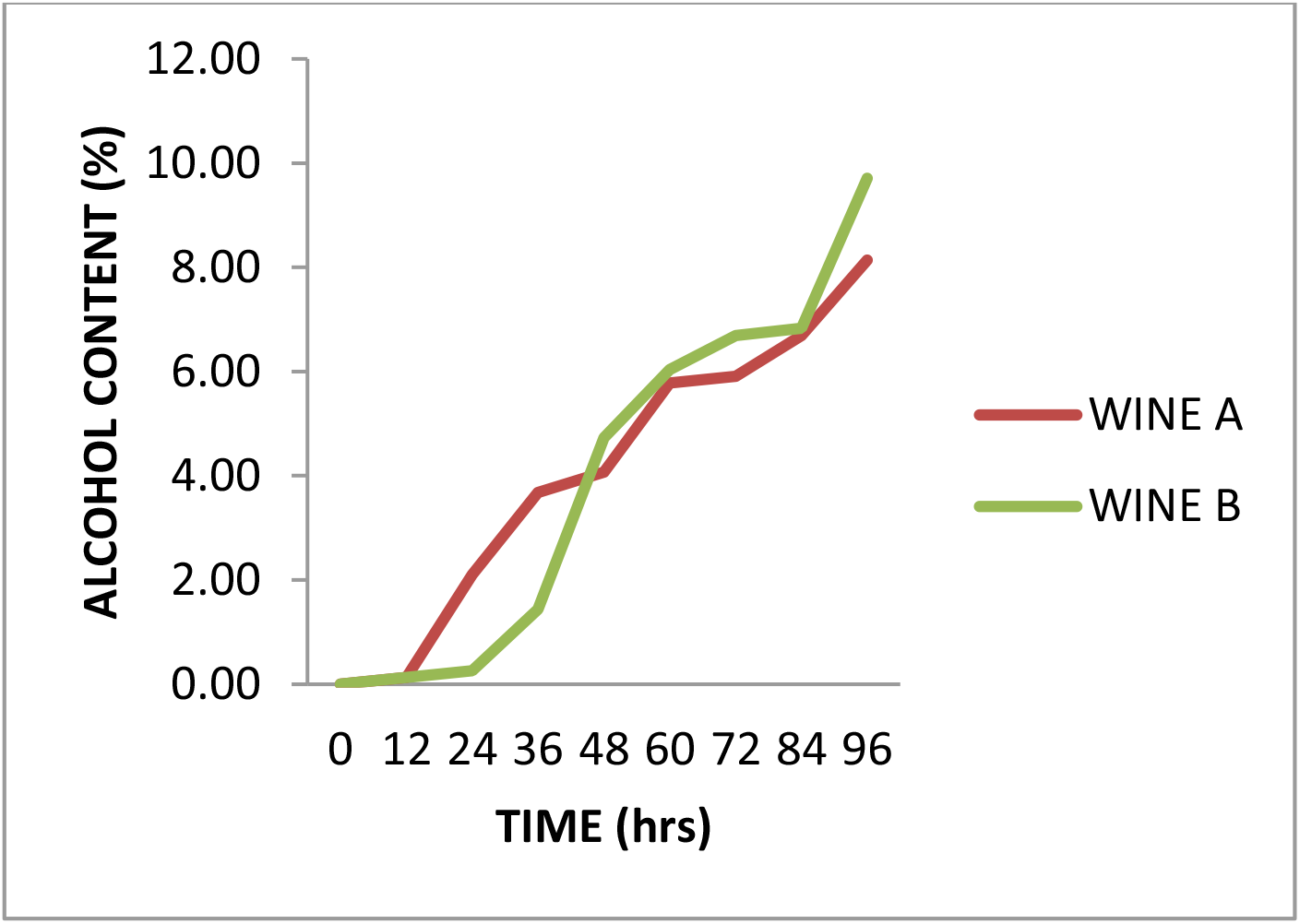
Variation in the alcohol content of the wines during primary fermentation

Figure 6 shows total acidity of the wines during primary fermentation ranging from 1.0% to 1.56% in Wine A and 1.0% to 1.62% in Wine B. Yeast count carried out during primary fermentation is shown in Table 1. The post-secondary fermentation analysis is indicated in Table 2. Proximate analysis of the wines as shown in Table 3, compared favourably after clarification and maturation. Sensory evaluation rated the acceptability of the wine at P>0.05 as portrayed in Table 4.

**Fig. vi:**
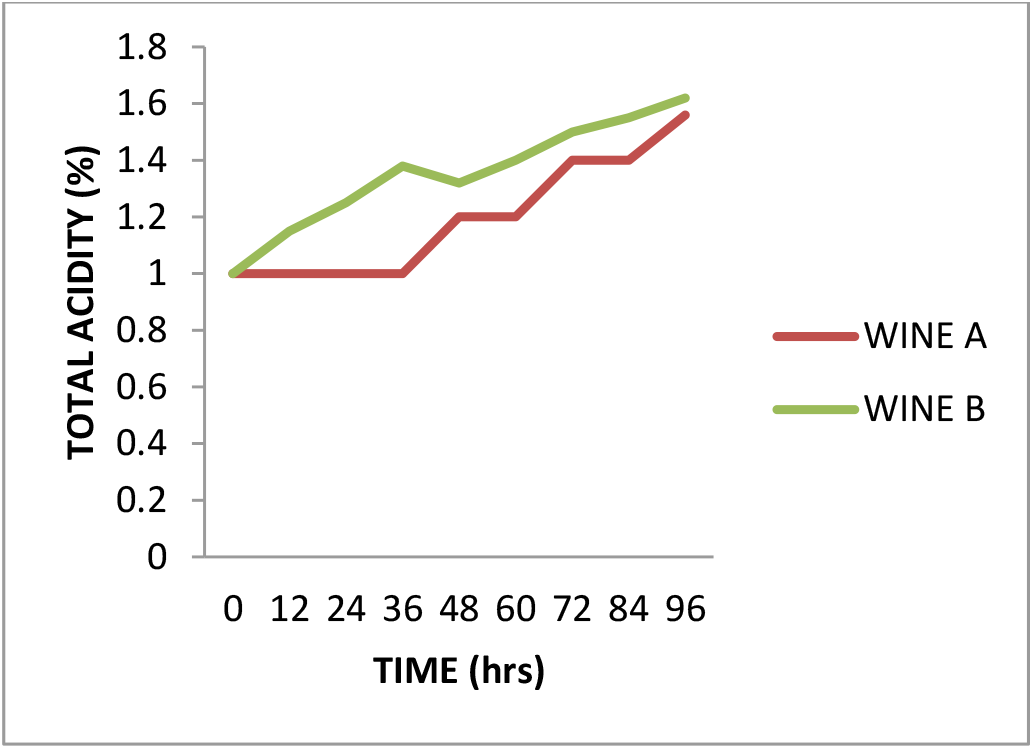
Variation in the total acidity of the wines during primary fermentation

**Table 1:**
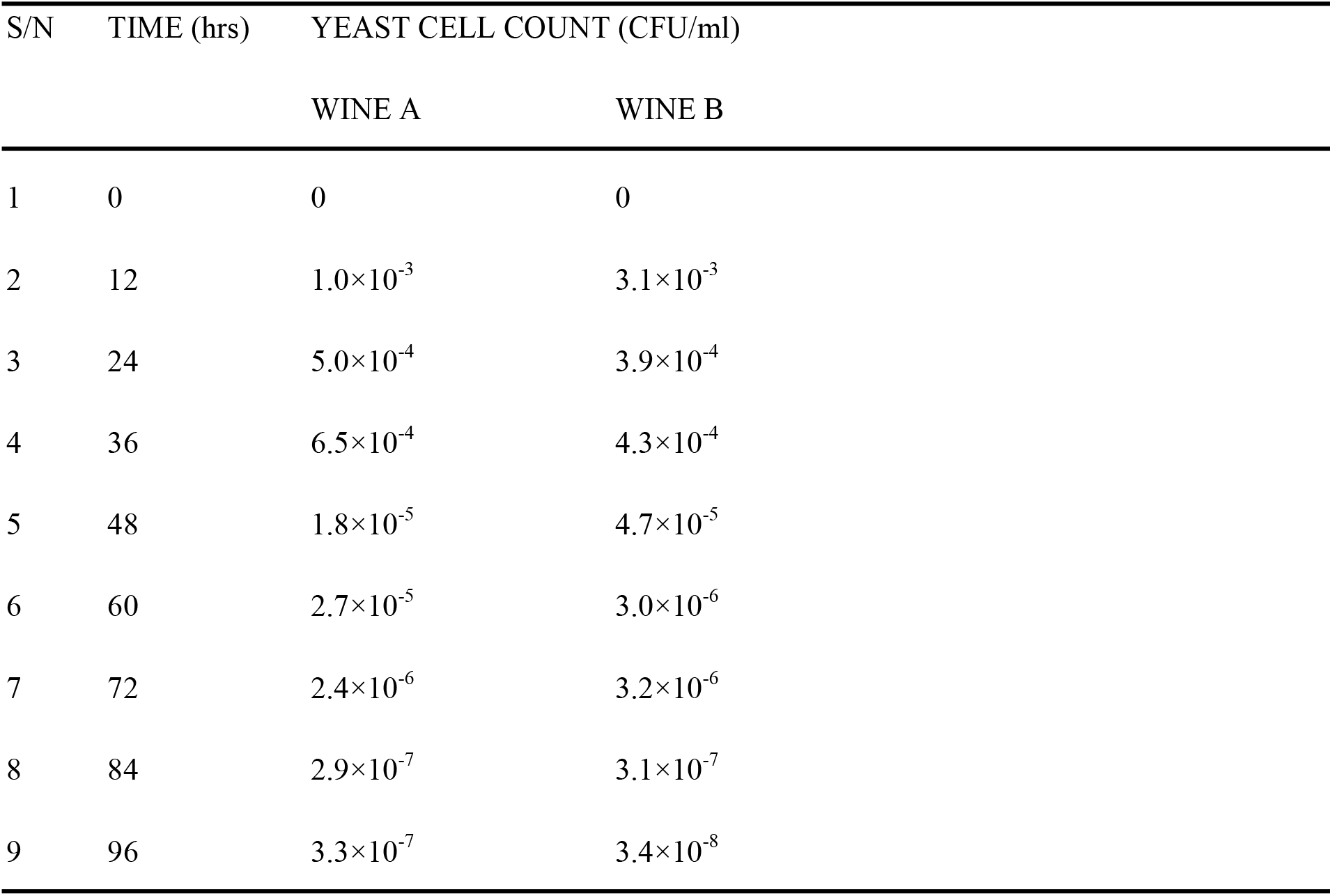
Yeast cell count.

| S/N | TIME (hrs) | YEAST CELL COUNT (CFU/ml) |  |
| --- | --- | --- | --- |
|  |  | WINE A | WINE B |
| 1 | 0 | 0 | 0 |
| 2 | 12 | $1.0 \times 10^{-3}$ | $3.1 \times 10^{-3}$ |
| 3 | 24 | $5.0 \times 10^{-4}$ | $3.9 \times 10^{-4}$ |
| 4 | 36 | $6.5 \times 10^{-4}$ | $4.3 \times 10^{-4}$ |
| 5 | 48 | $1.8 \times 10^{-5}$ | $4.7 \times 10^{-5}$ |
| 6 | 60 | $2.7 \times 10^{-5}$ | $3.0 \times 10^{-6}$ |
| 7 | 72 | $2.4 \times 10^{-6}$ | $3.2 \times 10^{-6}$ |
| 8 | 84 | $2.9 \times 10^{-7}$ | $3.1 \times 10^{-7}$ |
| 9 | 96 | $3.3 \times 10^{-7}$ | $3.4 \times 10^{-8}$ |

**Table 2:** Analysis of wine after secondary fermentation.

| S/N | Parameters | Wines |  |
| --- | --- | --- | --- |
|  |  | WINE A | WINE B |
| 1 | Volume of wine | 439ml | 240ml |
| 2 | pH | 2.29±0.1 | 3.25±0.2 |
| 3 | Specific gravity (kg/l) | 1.021±0.02 | 1.027±0.03 |
| 4 | Temperature (°C) | 27.00±1.1 | 26.50±1.1 |
| 5 | Total Acidity (%) | 1.57±0.2 | 2.11±0.1 |
| 6 | Alcohol Content (%) | 8.40±2.9 | 9.84±3.6 |

Values are expressed as mean ± standard deviation

**Table 3:**
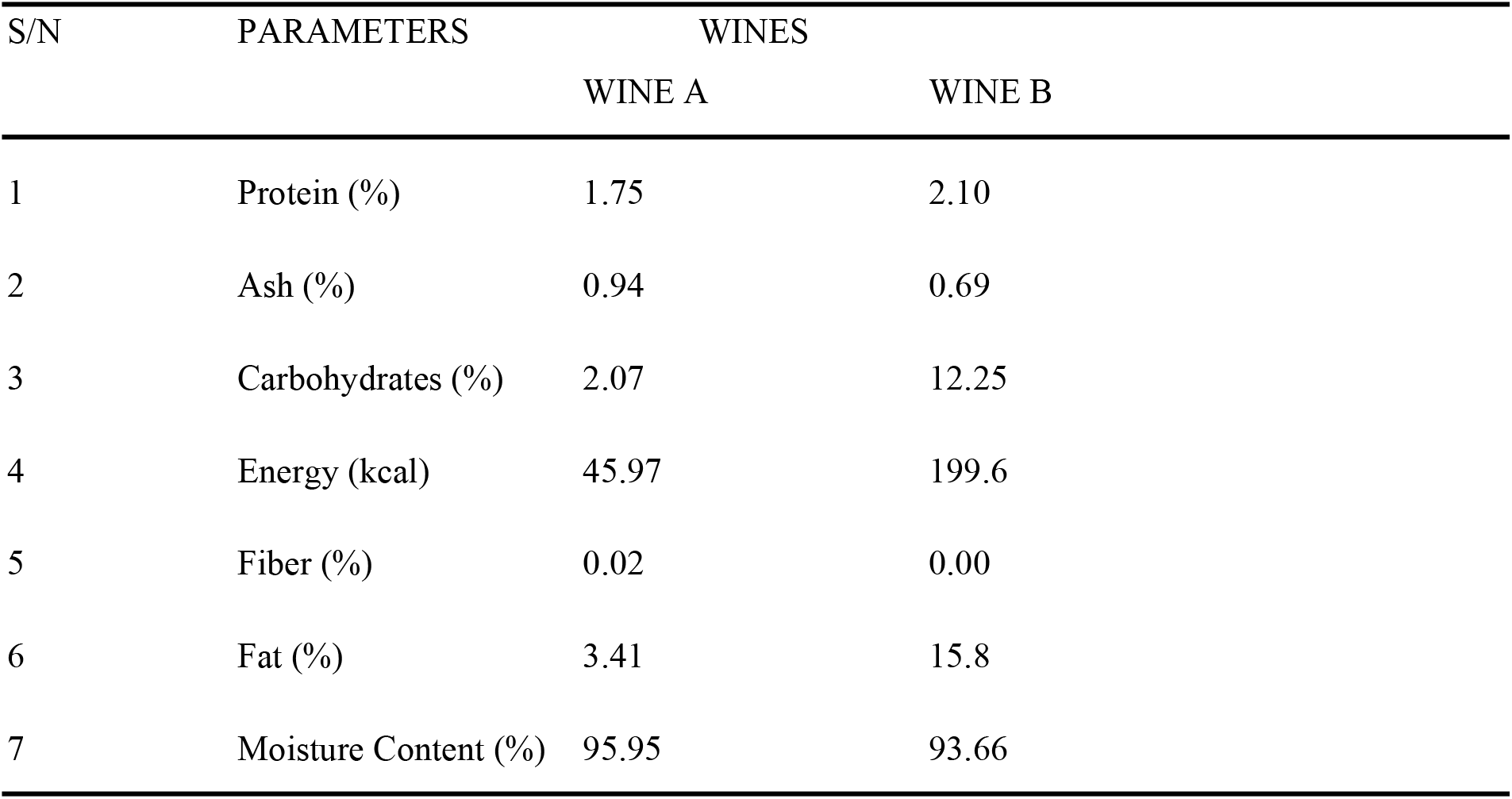
Proximate analysis of the final wine.

**Table 4:** Sensory evaluation of the wine.

| S/N | PARAMETERS | WINES |  | P value |
| --- | --- | --- | --- | --- |
|  |  | WINE A | WINE B |  |
| 1 | Color | 3.8 | 4.0 | >0.05 |
| 2 | Aroma | 4.2 | 3.8 | >0.05 |
| 3 | Clarity | 3.2 | 2.6 | >0.05 |
| 4 | Taste | 3.0 | 3.4 | >0.05 |
| 5 | Flavor | 3.0 | 3.8 | >0.05 |
| 6 | Overall acceptability | 3.4 | 3.5 | >0.05 |

Significant difference is taken at $P < 0.05$

## Discussion

The pH values of watermelon and pineapple wine (Wine A) and watermelon and banana wine (Wine B) ranged from 3.69 to 3.32 during primary fermentation, thereby maintaining an acidic range and conferring stability on the wine sample. As shown in figure 1, there was an observed continual decrease in the pH level as the hours decreased throughout the period of primary fermentation, revealing that the pH of this present study compares favorably with the reports of Nzabuheraheza and Nyiramugwera (2014) who produced golden wine from passion fruit, mango and pineapple, Ogodo *et al*. (2015) for mixed fruit (pawpaw, banana and watermelon) wines, Sahu *et al*. (2012) for tendu wine and Okeke *et al*. (2015) for mixed fruits (pineapple and watermelon) wine. Agbor *et al*. (2011) noted that maximum pH for mixed fruit wine is 3.5. As fermentation progressed there was a continuous reduction in pH values of the fruit wines, the changes observed with the pH in the wine could be due to the acid production during the fermentation by the microorganism (Idise and Odum, 2011). According to Won *et al*. (2015), the fermentation process which has a pH lower than 4.0 is regarded as homofermentative, devoid of contamination. Based on this assertion, the mixed fruit wine fermented using *Saccharomyces cerevisiae* isolated from palm wine is considered safe for human consumption. According to Chilaka *et al*. (2010), during the fermentation process, low pH inhibits the growth of spoilage organisms and rather makes the environment favorable for the growth of desirable microorganisms. Therefore, the final products, watermelon and pineapple (Wine A), watermelon and banana (Wine B) mixed fruit wines with pH of 2.29±0.1 & 3.25±0.2 respectively, could be considered safe for consumption due to the absence of undesirable microorganisms. Wine makers use pH as a way to measure ripeness in relation to acidity as low pH wines will taste tart and crisp while high pH wine will be more susceptible to bacterial growth as reported by (Vinny, 2009), pH is important in wine production because it improves the chemical and biological stability of the wine.

The change in temperature during fruit wine production presented in figure 2 shows a fluctuating temperature range of 30°C to 27.5°C from 0hrs to 96hrs in both mixed fruit wines with a final range of 26.50 ± 1.1°C in Wine A and 27.00 ± 1.1°C in Wine B; this is as a result of the fermentation activity and ambient temperature changes. This increase in temperature at 0hr shows that the higher the temperature the faster the yeast is able to convert the must from sugars to alcohol and it is seen that from 72hrs to 96hrs there was a slight decrease in temperature which steadied for a period of time showing that as the hours increase the sugar content and temperature decreases, these observed changes in temperature of the wine could be due to microbial succession arising from microbial metabolic activities as reported by Okafor (2007). Similar observations were recorded for apple mango wine produced by Musyimi *et al*. (2013), banana wine produced by Sevda *et al*. (2011) and mixed fruit (watermelon, banana and pawpaw) wine by Ogodo *et al*. (2015). The temperature of fermentation is usually from 25°C to 30°C, this makes yeast an important microorganism for fermentation. The type of fruit used also affects the temperature as various tropical fruits require different fermenting temperatures. The ideal temperature for fermentation of fruit wines is typically between 20°C to 30°C, if the temperature of the fermenting wine drops below this range, the fermentation process may slow down or stop altogether, while temperature above this range can lead to off-flavors and unwanted aromas.

In this study, the reducing sugars as shown in figure 3 for Wine A shows a decrease in value from 0.0440 to 0.0230mg/1000ml and 0.0448 to 0.0340mg/1000ml in Wine B as the number of hours increases from 12hrs-96hrs. This reduction is due to the conversion of sugars to alcohol during fermentation, this decrease in the reducing sugar implies that the wine produced is good for consumption as this lowers the risk of developing overweight, obesity and diabetes. Examples of these reducing sugar includes; galactose, glyceraldehyde, glucose, fructose, ribose and xylose which means a reduction in values for reducing sugar in the wine also brings about low carbohydrate in the wine as reported by Zoecklin *et al*. (1990). The mixed fruit wine compares similarly to reports of Enrika *et al*. (2018) for pineapple and banana wine, Kotecha *et al*. (1994) for banana wine with a reducing sugar of 0.044 ± 0.002%. In a related study, Umeh *et al*. (2015) reported that during fermentation of pawpaw wine must using *Saccharomyces cerevisiae* isolated from burukutu, the reducing sugar content steadily decreased from 16.70 to 1.10% between Days 0-14. Although reducing sugar content of the pawpaw wine was higher compared with wine produced using *Saccharomyces cerevisiae* isolated from palm wine, both results showed the same trend.

In the case of specific gravity of the fruit wines produced in this study as shown in figure 4, the specific gravity of Wine A and Wine B gradually decreased in values as observed throughout the period of fermentation, it reduced as fermentation hours of the wine increased ranging from 1.085 to 1.023kg/l during primary fermentation to a final range of 1.021±0.02kg/l and 1.027±0.03kg/ in Wines A and B respectively, after 21 days of secondary fermentation, this steady decrease is due to the activities of the yeast which feeds on the sugar of wine into alcohol and carbon dioxide through the process of fermentation because all the sugars present in the starting fermentation has been used by the yeast in the fermentation reaction to produce carbon dioxide and alcohol. *Saccharomyces cerevisiae* isolated from palm wine has been reported to reduce specific gravity of fruits wines as reported by Okeke *et al*. (2015). This is similar to the reports of Idise and Odum (2011) for banana wine, Yusufu *et al*. (2018) for watermelon juice and ginger extract, Nzabuheraheza and Nyiramugwera (2014) for golden wine, but higher as compared to Ogodo *et al*. (2015) for mixed fruit (watermelon, banana and pawpaw) wine and Reddy and Reddy (2005) for mango wine. In a related study, Chilaka *et al*. (2010) reported that there was a gradual decrease in the specific gravity of passion fruit wine, watermelon fruit wine and pineapple fruit wine during fermentation of the musts, though in agreement with the trend observed in this study, they had lower values which ranged from 0.91 to 0.92, 0.90 to 0.91 and 0.94 to 0.96 kgm^-3^, respectively.

A steady increase in alcohol content was observed in the fruit wine throughout the period of primary fermentation and this is shown in figure 5, the concentration of alcohol in the fruit wines were observed to range from 0 to 9.71% during primary fermentation. According to Clement *et al*. (2005), high alcohols are known to be important precursors for the formation of esters, which are associated with pleasant aromas, hence, the importance of alcohol content in wine is to provide balance in the taste, texture, aroma and quality of the wine, thus, the fruit wines in this study have a high percentage of alcohol thereby giving it a good flavor, aroma and taste. The type of fruits used and ripening levels were deliberate because it causes an increase in the alcohol content of the fruit wines. The alcohol content of the fermenting wine increased during fermentation and this can be attributed to yeast metabolism by continuous utilization of the sugar content. Ethanol produced leads to an increase in the alcohol content of the fermenting must and this continues until the available sugar in the fermenting must is utilized. This corresponds to the reports of Musyimi *et al*. (2013) who prepared wine from an apple mango variety with an alcohol content of 9.44%, Won *et al*. (2015) for apple wine mash with an alcohol content of 11.97%, Chilaka *et al*. (2010) who produced passion, watermelon and pineapple wines with an alcohol content of 12.42%, 10.44% and 12.80%, respectively. The final alcohol content was 8.40±2.9% in Wine A and 9.84±3.6% in Wine B. This observation does not correspond with the reports of Ogodo *et al*. (2015) for mixed fruit wines and

Nzabuheraheza and Nyiramugwera (2014) for golden wine which had higher alcohol contents. In terms of alcohol content for table wine, the European Economic Community (EEC) recommends that it should be within the range of 8.5% to 19.5% (Umeh *et al*., 2015). Interestingly, the watermelon and banana wine produced using *Saccharomyces cerevisiae* isolated from palm wine met the alcohol content requirement of EEC which qualifies the product as a good table wine and furthermore agrees with the findings of Sandipan and Subhajit (2011) that wines with 7-14% of alcohol are considered as table wines.

Acidity plays a vital role in determining wine quality by aiding the fermentation process and enhancing the overall characteristics and balance of the wine. Lack of acidity will mean a poor fermentation (Berry, 2000). It was observed that the total acidity of the mixed fruit wines consistently increased from 1.0% to 1.62% as shown in figure 6 therefore, leading to a range of 1.57±0.2% in Wine A and 2.11±0.1% in Wine B after secondary fermentation. This indicates an increase in the total available hydrogen ions present in the must, also, an increase in total acidity is an indication of the accumulation of tartaric acid level due to yeast metabolism and the organism that produces the enzyme that allows glucose to be fermented have adapted to the acidic conditions (Awe and Nnadoze, 2015). This corresponds with the reports of Okeke *et al*. (2015) who produced pineapple and watermelon wine (1.01%), Ray *et al*. (2011) for purple sweet potato wine (1.35g/100ml) and Panda *et al*. (2014b) for bael wine (2.05g/100ml) and differs higher with reports of Ogodo *et al*. (2015) for mixed fruit wines (0.35 to 0.88%) and Yusufu *et al*. (2018) for watermelon juice and ginger extract (0.04 to 0.10%). The changes in the total acidity of the wine produced within the period of fermentation shows the occurrence of Malo-lactic fermentation. Malo-lactic fermentation arises from the succession of the yeast cells by lactic acid bacteria after 48hrs of fermentation. The presence of a malo-lactic fermentation is a desirable phenomenon in wine production due to the attendant buttery flavor (Idise, 2011).

From Table 1, the yeast cell count of both wines is observed to increase, indicating the presence of sugar in the must that can be utilised for growth purposes. At the initial, culturable microorganisms were not detected in the mixed fruit wine. Between 12 hours and 96 hours, the result indicates that there was an increase in the mean count of the colonies in the fermenting mixed fruit musts. In a related study, Zainab *et al*. (2018) reported a rapid increase in the population of the yeast cells involved in the fermentation of watermelon wine within 54h from the time the process commenced. The increase in the population of the yeast is an indication that an adequate amount of sugar was available for the organism to utilize. During the course of this experiment, the fermentation yeast, *Saccharomyces cerevisiae* was the only organism isolated from the mixed fruit wine as a result of the multiplication of the yeast present in the fermenting must and the addition of yeast nutrients. This is an indication that the wine is of good quality and safe for consumption. This observation may be due to the addition of sodium metabisulphite which serves as a sterilizer and inhibits the growth of unwanted organisms if used in appropriate amounts and monitored properly, also, low pH values, high acidity and high alcohol contents of the wines which are known to inhibit the growth of pathogens and gives fermenting yeast a competitive advantage in natural environment as reported by Reddy and Reddy (2005) and Chilaka *et al*. (2010). The presence of the growth of the yeast in the wine could be due to the average alcoholic content which did not exceed the ethanolic tolerance level of the yeast used for fermentation. The rapid multiplication of the yeast could be attributed to the addition of potassium phosphate which helps to ensure that the yeast has access to sufficient phosphorus which is important for yeast growth and multiplication, as well as for the production of certain fermentation byproducts. The addition of ammonium sulfate provides essential nutrients especially nitrogen for microbial growth, as well as its buffering capacity and affordability, the addition of magnesium sulfate serves as a source of magnesium and sulfate ions. Magnesium is an important nutrient for yeast during fermentation, while sulfate ions contribute to the flavor and aroma of the finished wine.

The proximate composition results in Table 3 shows a high moisture content of 95.95% in Wine A and 93.66% in Wine B. The moisture content of a sample determines how shelf stable a product will be and the overall nutritional value of the sample, therefore proximate analysis carried out on the mixed fruit wines confirmed that the wine is good for healthy consumption and the high moisture content accounts for the perishable nature of the fruits and their short life under normal conditions. High moisture content makes beverages suitable as a refreshing and quench-thirsting product which is a characteristic of a good beverage, similarly agreeing with the report of Okeke *et al*. (2015). Ash content of 0.94% was observed in Wine A and 0.69% in Wine B, in a similar study by Kantiyok *et al*. (2021), ash content was reported to have a value of 0.51% for pawpaw and watermelon wine. Fat content of 3.41% was observed in Wine A and 15.8% in Wine B, according to Kinsella (1976), fat absorption can be influenced by the lipophilicity of protein. Crude fiber of 0.02% was noted in Wine A and was observed to be 0.00% in Wine B, the absence of crude fiber further demonstrates the desirable nutritive quality of the fruit wines produced (Kantiyok *et al*., 2021). Protein content of 1.75% in Wine A and 2.1% in Wine B was observed and this may be due to the higher protein present in the proportion of banana fruit used. The protein content differs comparably from red wine, pawpaw wine and banana wine which from the report of Awe *et al*. (2013) has a lower value from the mixed fruits (watermelon, pineapple and watermelon) wine produced. In Wine A, carbohydrates of 2.07% and energy value of 45.97kcal was observed, Wine B also contained a reasonable amount of carbohydrates with a value of 12.25% and energy value of 199.6kcal, which invariably accounts for their high caloric value suggesting the presence of energy source for metabolic activity of the yeast (Kantiyok *et al*., 2021).

The sensory evaluation conducted proves that the wines produced from the mixtures of watermelon & pineapple and watermelon & banana can be considered to be a good quality wine with an acceptance rate of 3.4 and 3.5 respectively, as shown in Table 4, making up to 80%. The wine sample shows that wine A is clear with a pale red color while wine B is clear with a pale-yellow color, the wine’s clarity can be ascribed to the addition of gelatin in the wine after secondary fermentation. These wines were reported by the assessors to be slightly alcoholic with a fruity-like flavor. The type of aroma produced during wine making was dependent on the yeast, environmental factors and physiochemical characteristics of the ‘must’. In texture, the wines produced were completely watery at the end of fermentation with bubbles formed on the surface. This result is similar to that recorded by Amerine *et al*. (1980), who stated that as fermentation rate proceeded, gas was formed and this rose through the liquid during active fermentation, froth or foam is formed on the surface. The overall acceptability compares favorably with the reports Ogodo *et al*. (2015) for mixed fruit wines, Sahu *et al*. (2012) for tendu wine and Panda *et al*. (2014a) for sapota fruit wine. The sensory evaluation of the wines in this study do not differ significantly (P>0.05) and its acceptability rated as watermelon and banana wine > watermelon and pineapple wine. Generally, this study has revealed the effectiveness of palm wine yeast strain in wine production from the test fruits (watermelon, banana and pineapple).

## Conclusion

This study has demonstrated that wine of good quality could be produced from indigenous fruits such as watermelon, banana and pineapple as substrates for wine production and the effectiveness of *Saccharomyces cerevisiae* isolated from local alcohol beverages like palm wine for mixed fruit wine fermentation has proven to be an efficient substrate for wine production. Fermenting fruits into wines not only provides different varieties of wine products but also confers nutritive values to consumers, improves the shelf life of perishable fruits and reduces waste. This study has given a broad insight on the use of *Saccharomyces cerevisiae* from other sources such as locally fermented beverages to ensure advancements in fruit wines which could further lead to better aroma, color, flavor, taste and bioactive components which offer numerous health benefits that would in turn lead to a wider commercialization of the product thereby contributing to the economy, encourage investments and long-term capital to consumers.

## Acknowledgements

The authors wish to acknowledge the efforts of Mr Igiri, I. I. in wine analysis and the panel of wine judges in ascertaining the acceptability of the wines.

